# Concomitant immunity to M. tuberculosis infection

**DOI:** 10.1101/2022.08.02.502562

**Authors:** Louis R. Joslyn, JoAnne L. Flynn, Denise E. Kirschner, Jennifer J. Linderman

## Abstract

Some persistent infections provide a level of immunity that protects against reinfection with the same pathogen, a process referred to as concomitant immunity. To explore the phenomenon of concomitant immunity during *Mycobacterium tuberculosis* infection, we utilized *HostSim*, a previously published virtual host model of the immune response following Mtb infection. By simulating reinfection scenarios and comparing with data from non-human primate studies, we predict that the durability of a concomitant immune response against Mtb is intrinsically tied to levels of tissue resident memory T cells (Trms) during primary infection, with a secondary but important role for circulating Mtb-specific T cells. Further, we compare *HostSim* reinfection experiments to observational TB studies from the pre-antibiotic era to predict that the upper bound of the lifespan of resident memory T cells in human lung tissue is likely 2-3 years. To the authors’ knowledge, this is the first estimate of resident memory T-cell lifespan in humans. Our findings are a first step towards demonstrating the important role of Trms in preventing disease and suggest that the induction of lung Trms is likely critical for vaccine success.

## Introduction

The human immune system has the potential to respond effectively to infection with many pathogens. After a first encounter with a pathogen (primary infection) is cleared, immunological memory is typically protective against a second infection event (reinfection) at a later date. Some persistent pathogens (i.e. those that are not cleared) also confer protection against secondary infection (1–5). This latter phenomenon, known as *concomitant immunity*, was first described in Stedman’s medical dictionary as “infection-immunity” (6). Formally, concomitant immunity is “the paradoxical immune status in which resistance to reinfection coincides with the persistence of the original infection” (7,8).

Concomitant immunity is thought to be pathogen-dependent and thereby mediated by different immune modalities depending on infection type. For example, persistent infection with lymphocytic choriomeningitis virus (LCMV) promotes a greater accumulation of relatively long-lived effector-like memory T cells at nonlymphoid sites (9). In *Leishmania major* infection, relatively short-lived CD4+ regulatory T cells play a role in concomitant immunity by preventing total clearance of the pathogen during persistent infection (7) and it was recently found that if CD4+ effector T cells are not sustained, concomitant immunity against reinfection was inadequate (10). Across multiple types of parasitic infections, separate and distinct cellular and antibody immune responses develop that are consistent with acquisition of concomitant immunity (1–5).

In tuberculosis (TB), which remains a significant global health crisis that was exacerbated by the COVID-19 pandemic, it is unclear if persistent *M. tuberculosis* (Mtb) infection in humans confers enduring protection against reinfection. The hallmark of TB, a primarily pulmonary disease, is the formation of lung granulomas: organized immune cellular structures that surround Mtb (11). Granulomas are composed of various immune cells, including macrophages and T cells (primarily CD4+ and CD8+ T cells, although other unconventional T cell phenotypes and B cells are also present, reviewed in (12)). T cells have well-established critical functions against Mtb (13–17), but unlike other infections, T cells arrive approximately one month after primary infection (18). The delays in lymph node T cell priming, activation, and trafficking through blood to lungs is characteristic of Mtb infection (19,20) and may be due to the slow growth of the bacilli and limited ‘danger’ signals (21) at early stages of infection during Mtb growth in the lungs (18).

Concomitant immunity against Mtb appears to provide at least some protection against reinfection. Observational cohort studies from natural infection case studies suggest that individuals with latent TB (LTBI – i.e., those who are persistently infected but are clinically asymptomatic) had a 35 - 80% lower risk of progression to active TB after re-exposure to Mtb compared to uninfected individuals (22–26). In those studies, it was not possible to determine whether actual reinfection occurs, since active TB was the only outcome measure. Nonetheless, in a seminal study of nursing and medical students, those who were tuberculin skin test positive (TST+) and had LTBI were less likely to develop active TB during their years in hospital training than those who were TST negative (22). Further, other studies in mice suggest that concomitant immunity is not fully protective against secondary infection with Mtb, but that bacterial burden from a secondary infection is reduced when compared to primary infection (27,28). Cadena et al. showed that in non-human primates (NHP) concomitant immunity was robust against a secondary infection with Mtb (29). However, due to the inherent constraints of NHP studies, Cadena et al. were unable to conduct re-exposures at multiple time points after primary infection and therefore were unable to quantify the potential longevity of a concomitant immune response. Based on the observation that Mtb-specific T cells reside in uninvolved lung tissue prior to reinfection, Cadena et al. hypothesized that resident memory T cells prevent the establishment of reinfection.

The defining feature of Trms is their persistent residence within nonlymphoid tissues and inability to circulate through blood stream and lymphatics (30). These cells act as a sentinel against future infection and have been shown to be protective against infections with influenza virus, herpes simplex virus, and human immunodeficiency virus (reviewed in (30)). Further, skin Trm cells have been shown to provide concomitant immunity in the case of cutaneous leishmaniasis (31). Relatively few studies have examined Trms in the context of TB (32). However, mucosal administration of the BCG vaccine in mice, as well as adoptive transfer of CD8+ Trms, have demonstrated enhanced protection against Mtb, presumably through the ability of Trm to respond quickly following infection (33). Additionally, intravenous administration of BCG in NHPs provided robust protection against infection, and BCG IV vaccinated NHP had high levels of Trm in lungs in contrast to intradermal or aerosol BCG vaccinated NHP (34). In general, Trms are thought to develop during the adaptive immune response to primary Mtb infection and have been identified within uninvolved lung tissue of infected hosts (29). However, experimental studies have so far been unable to define their role in concomitant immunity against Mtb and the longevity of this cell population in the lung has not been well-characterized in NHPs or humans.

As a complementary approach, mathematical and computational modeling can predict mechanism and timing of major immune events beyond the timeline of experimental studies. In TB, modeling has been used to explore various aspects of granuloma formation (35,36), drug-dynamics (37,38), and immune cell and cytokine dynamics within lung granulomas (36,39–41). The advantages of a modeling approach are well-suited to answer outstanding questions about reinfection and the potential longevity of Trm cell populations in the human lung.

We previously calibrated and validated *HostSim*, a whole-host modeling framework that captures key elements of TB immune responses and pathology across lungs, lymph nodes and blood (42), using multiple datasets derived from published NHP studies and have shown the ability of this model to capture heterogeneous host-scale clinical outcomes such as infection clearance, control (LTBI) or active disease. Here we used the *HostSim* framework to address two outstanding questions about Trms and concomitant immunity in TB: Do Trms mediate concomitant immunity in Mtb? Can we predict the lifespan of Trms in primate lung tissue and therefore the duration of concomitant immunity against Mtb infection?

## Results

### Resident Memory T cells (Trm) are the main mediators of concomitant immunity against Mtb reinfection

In NHPs, ongoing primary infection with Mtb confers protection against reinfection 16 weeks later (29). Four weeks after the secondary inoculation, animals were necropsied and granuloma bacterial burdens from both primary and secondary inoculation events were obtained. By sampling the parameter space of *HostSim*, we created a population of 50 virtual hosts and simulated the virtual hosts following the NHP experimental protocol (Figure 1A). We then compared *in silico* granuloma CFU levels to the NHP *in vivo* granuloma CFU levels from primary infection and reinfection.

**Figure 1:**
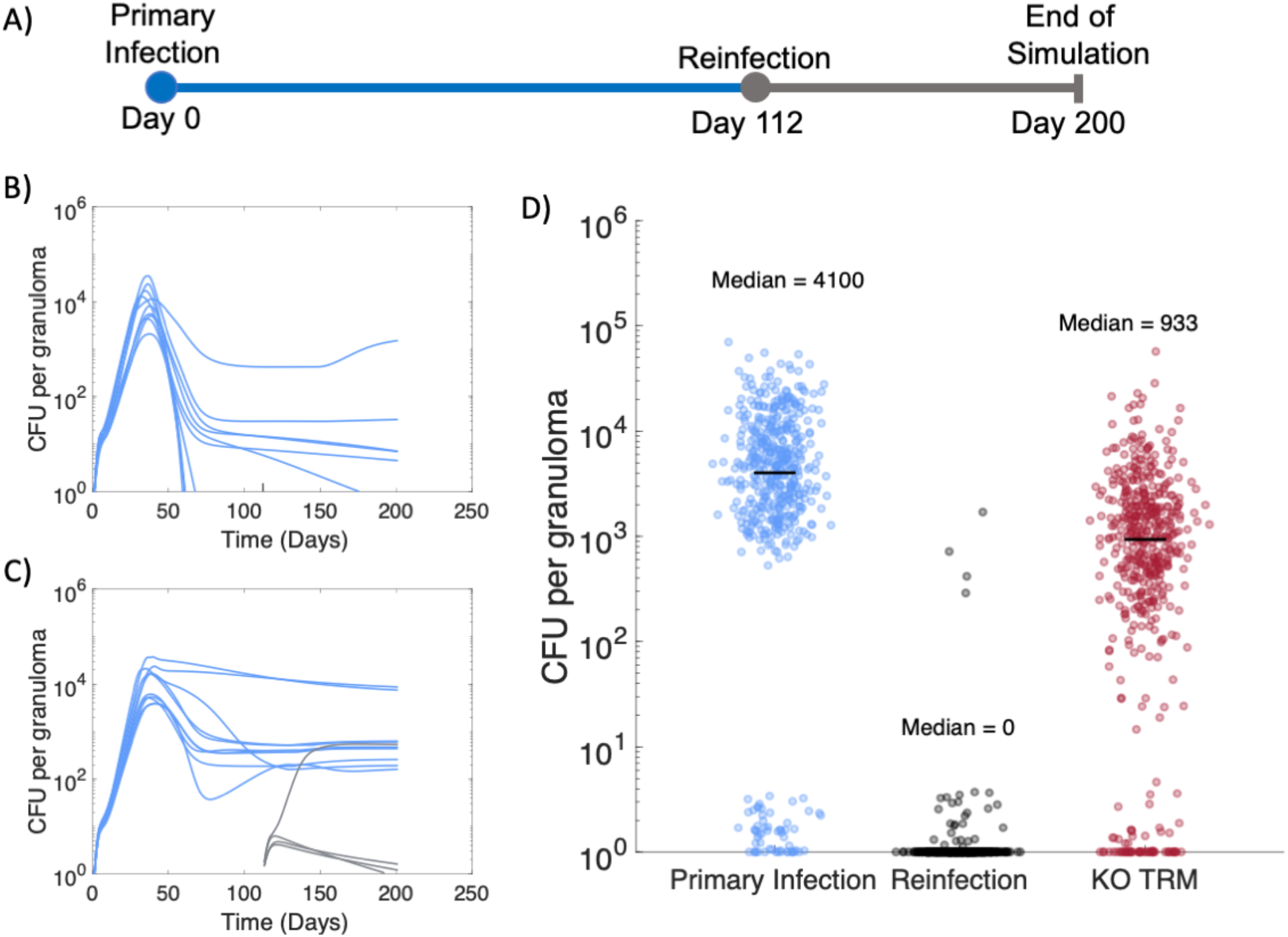
Resident memory T cells protect against establishment of reinfection granulomas in virtual population of 50 hosts. A) Experimental protocol for reinfection study. Primary infection (blue) with 10 CFU inoculum occurred at day 0 and reinfection (gray) with 10 CFU inoculum occurred at day 112, matching the NHP study protocol (29). Granuloma CFU trajectories from two representative hosts shown in panel (B) & (C). (D) Across 50 virtual hosts, reinfection granulomas exhibit greater sterilization compared to primary infection granulomas 28 days post-infection, as previously observed in NHPs (29). We re-simulated the same 50 hosts but “knocked-out” (KO) the Trm cell population at reinfection (red data points). CFU per granuloma between KO Trm virtual hosts and the primary infection granuloma CFU was significantly different (Vargha and Delaney’s A measure = 0.76).

We added Trms to our *HostSim* modeling framework due to their potential role in mediating protection against reinfection (see Methods). With the addition of Trms into *HostSim*, we observe that concurrent Mtb infection limits the establishment of reinfection granulomas, matching observed reinfection dynamics in NHPs (29). We predict that the vast majority of reinfection granulomas are sterilized prior to 28 days post-infection using *HostSim* (Figure 1). Our simulations resulted in virtual hosts where reinfection was not established, i.e. no granulomas due to secondary challenge were detected (Figure 1B) and virtual hosts where secondary challenge resulted in granulomas (Figure 1C) (gray lines indicated reinfection granuloma CFU). For 33 out of the 50 virtual hosts, reinfection was not established, as these hosts had total sterilization of all reinfection granulomas prior to day 28. This is consistent with the NHP study, which showed 5 of 8 monkeys had total sterilization of reinfection granulomas at this same time point. If we re-simulate the 50 virtual hosts but delete (knockout, KO) Trms, we no longer match the NHP dataset. Intriguingly, reinfection granulomas in the KO study still contained significantly lower bacterial burdens (~5 fold less) than primary infection granulomas (Figure 1D, red data points). This suggests that while Trms are the main drivers of a concomitant immune response, additional immune cells also likely affect the growth of bacteria within reinfection granulomas.

### Predicting the lifespan of Trms and durability of concomitant immunity

While concomitant immunity is protective against reinfection at 112 days in NHPs, the duration of a concomitant immune response over time is not yet known (29). However, prospective cohort studies from the pre-antibiotic era of TB treatment predict that individuals with latent Mtb infection (Tuberculin-skin test positive, asymptomatic, infected for an undetermined period of time, spanning years) had a 79% reduced risk of developing active TB following re-exposure to Mtb compared to uninfected individuals (22). These studies were observational, and it was not possible to determine the exact time of initial infection or re-exposure as the only outcome measure was active TB cases. However, these studies provide an opportunity for a case study: we can use *HostSim* to predict the lifespan of lung Trm – which our *in silico* knock-out experiment (Figure 1D) predicts as a major contributor to concomitant immunity.

We performed three sets of virtual reinfection studies using the same 500 virtual hosts (see Methods) to predict the lifespan of lung Trms and measure longevity of a concomitant immune response in *HostSim*. We alter the death rate of lung Trms between the three sets of reinfection studies to predict Trm lifespan in primates.

To estimate the lifespans of Trms during TB, we classify hosts across our virtual populations as active, latent or Mtb eliminators following reinfection to identify which Trm lifespan best aligns with the results of the human cohort study (22). To this end, we display the breakdown of active TB (dark blue), LTBI (green) and Mtb eliminator (yellow) cases (as defined by the total lung bacterial burden [Supplementary Figure S1]) at day 200 post-reinfection for the 500 virtual hosts across 21 reinfection studies (Figure 2). When the lifespan of Trms is set to previous estimates in mice proposed from Morris et al. (*d*_*Trm*_ =0.03 cells/day) (43), concomitant immunity wanes quickly (Figure 2A, D). In contrast, if the lifespan of Trms lasts for multiple decades (*d*_*Trm*_ =0.0001 cells/day; Figure 2C, F), all virtual hosts sterilize or control reinfection. Each of these Trm lifespans result in predictions that are inconsistent with the human cohort studies. Only when the lifespan of Trms is ~25x longer than originally estimated by Morris et al. (*d*_*Trm*_ =0.0012 cells/day, calculated based on allometric scaling between humans and mice), does the reduced risk of active TB align with that of the human cohort studies (Figure 2B, E). Across the 21 reinfection studies in Figure 2B, the average reduced risk of active TB is 77% following reinfection. This is a prediction that is consistent with reduced risk estimates in humans (~79%, Andrews et al. (22)). Thus, we predict that the lifespan of an individual Trm is approximately 2-3 years.

**Figure 2:**
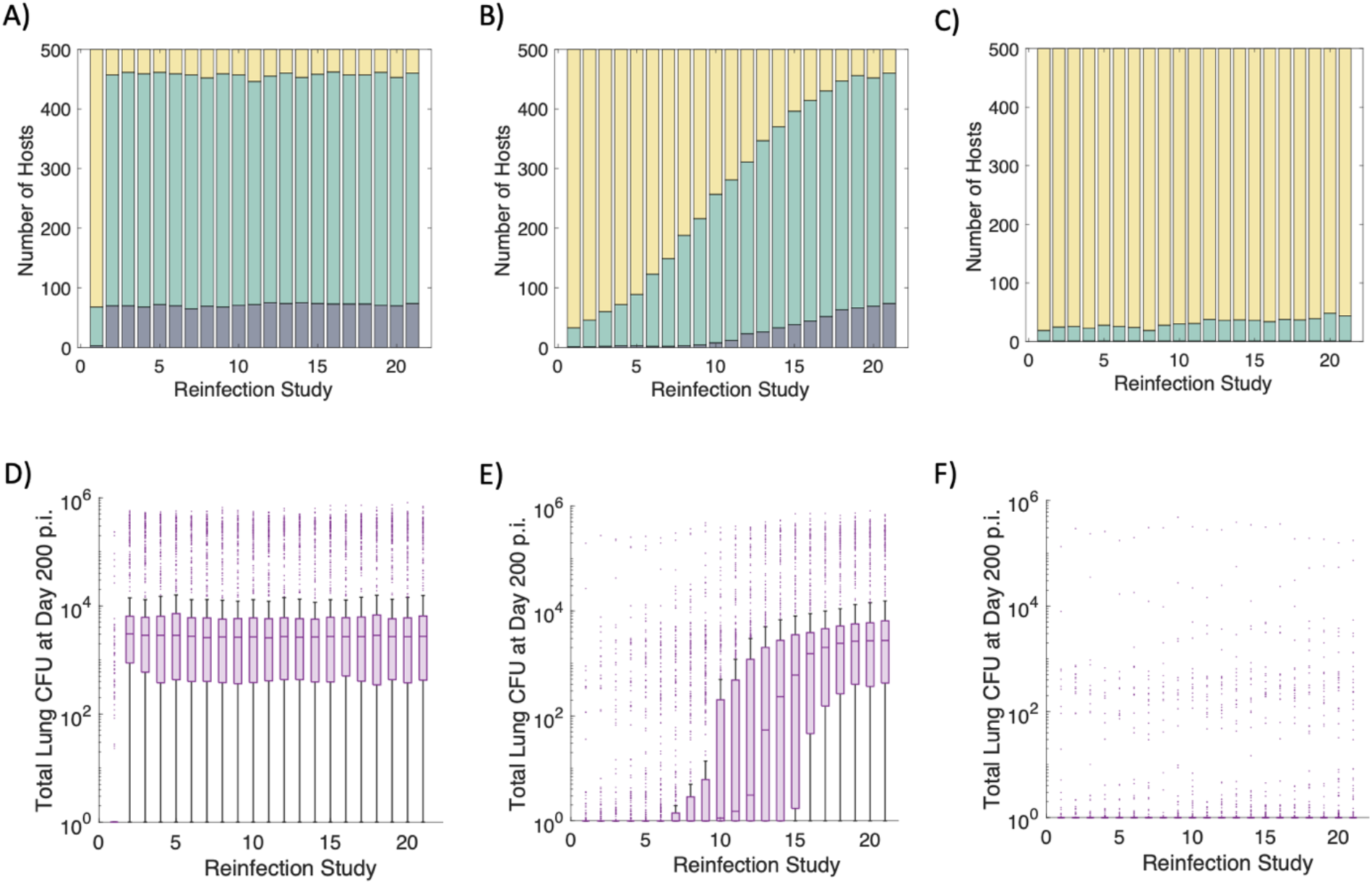
Clinical classifications and total lung CFU across three sets of virtual reinfection studies. Clinical classifications for 500 virtual hosts across 21 separate reinfection studies when the lifespan of Trms is varied from 33 days, 833 days, or 10,000 days (*d*_*Trm*_ = 0.03, 0.0012, or 0.0001 cells/day) (A, B, C, respectively). Each stacked bar chart shows the breakdown of active TB (dark blue), LTBI (green) or Mtb eliminator (yellow) hosts for each reinfection study. Clinical classifications are determined according to total lung CFU 200 days after reinfection. Box-and-whisker plots show the distribution of total lung CFU across the 500 virtual hosts for each reinfection study when *d*_*Trm*_ = 0.03, 0.0012, or 0.0001 cells/day (D, E, F, respectively).

### Mtb-specific blood T cells from primary infection offer protection against active TB during reinfection in absence of Trm populations

We showed that concomitant immunity is intrinsically associated with Trm lifespan (Figures 1 & 2). Intriguingly, at reinfection timepoints after the majority of Trm cells have died, we still note an average reduction of ~35% in active TB from reinfection compared to controls who were only infected once (Figure 2A, reinfection studies 2-21). We use these reinfection studies to identify mechanisms driving active TB following reinfection events that may occur after waning of Trm populations.

In Figure 3, we focus on the first set of reinfection studies, when the life span of Trms is 33 days (*d*_*Trm*_ = 0.03 cells/day), although the results herein are consistent across the three sets of reinfection studies (Supplementary Figure S3). In the first set of reinfection studies, the Trm population is no longer present by reinfection timepoint 212 days after primary infection (Figure 3A). For reinfection study numbers 2-21, we observed that the majority of virtual hosts still control infection, and are classified as LTBI, even without the presence of Trm populations in the lung (Figure 3A). Further, we observed that virtual hosts who control reinfection had higher counts of Mtb-specific effector (Figure 3B), effector memory (Figure 3C), and central memory (Figure 3D) T cells one day prior to reinfection as compared to hosts that went on to develop active TB after reinfection. Thus, we predict that the numbers of Mtb-specific T cells in the blood prior to reinfection is a key factor for protection against active TB following reinfection in the absence of Trms.

**Figure 3:**
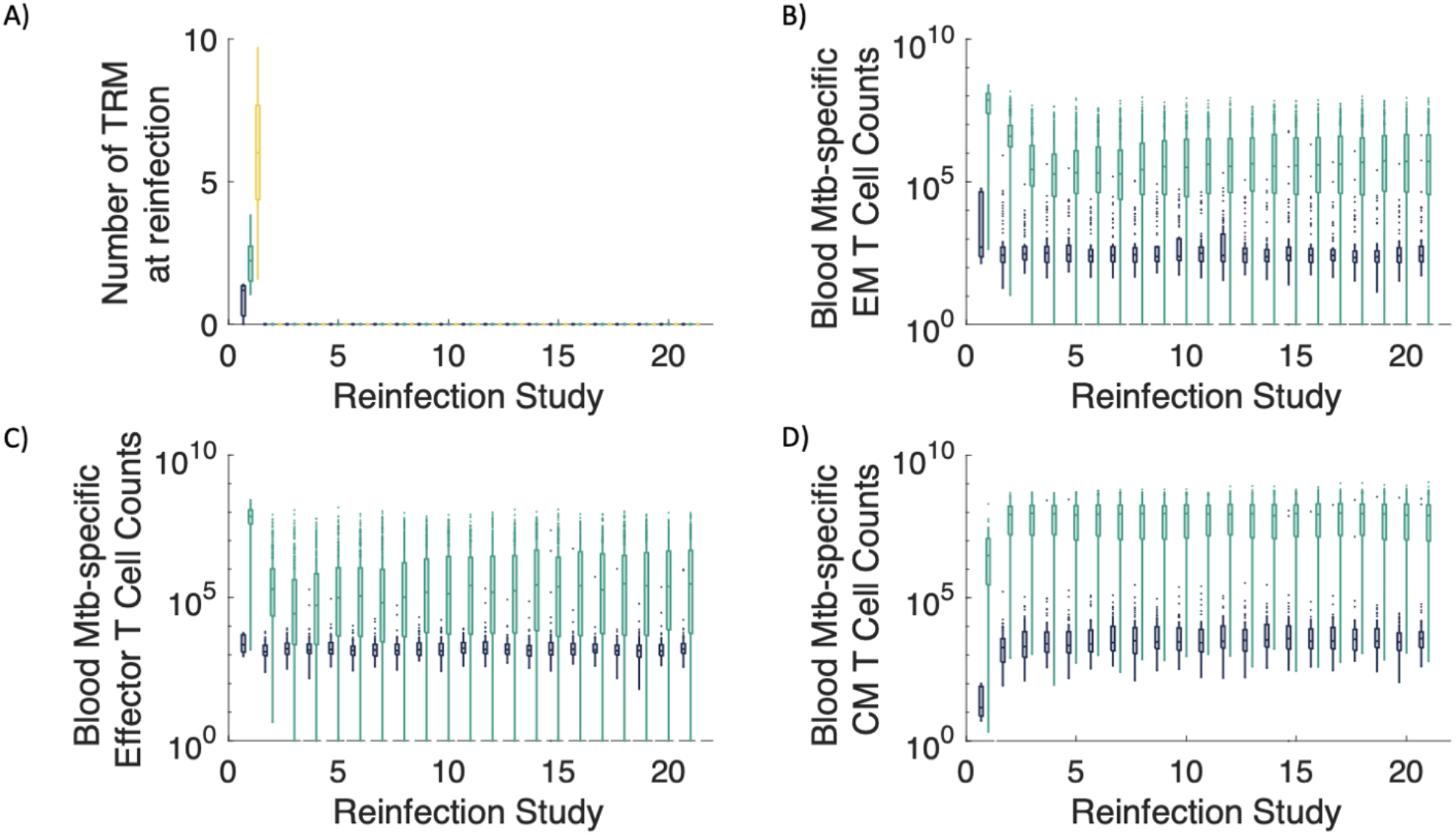
In the absence of Trms, Mtb-specific T-cell counts in blood delineate active vs. LTBI outcomes following reinfection. (A) In our first set of reinfection studies, we set the lifespan of Trms to be 33 days (*d*_*Trm*_ = 0.03 cells/day), and the Trm population wanes by day 212, the reinfection time point of reinfection study 2. Box-and-whisker plots show the distribution of Mtb-specific effector memory (EM), effector, and central memory (CM) T cell counts in the blood one day prior to the timepoint of reinfection for each reinfection study (B, C, D, respectively). We stratify the T-cell numbers according to a host’s total lung CFU at a time point 200 days later. If a virtual host was classified as an active TB case (total lung CFU>10^5^), the box and whisker plot is dark blue. If the virtual host was classified as an LTBI case (total lung CFU<10^5^), the box and whisker plot is green. Note that the vast majority of LTBI virtual hosts have larger numbers of Mtb-specific blood effector, effector memory and central memory T cells than that of their active TB case counterparts.

## Discussion

Concomitant immunity is a special case of immune memory that is generated when a host is re-exposed while they are currently harboring a primary infection by the same pathogen. However, concomitant immunity does not provide a robust and enduring immune response against reinfection for all persistent pathogens (44). For example, during chronic HIV infection, reinfection has been shown to clinically arise as early as one-year after primary infection (45). If a concomitant immune response is observed, then the cellular-mediators and longevity of immunity appear to be pathogen-dependent, ranging from memory T cell mediated immunity in chronic LCMV to extremely short-lived immunity in parasitic infection (7,9,10). In TB, concomitant immunity against reinfection appears protective, but the longevity and cellular mediators of such a response are not yet known.

To address these uncertainties, we utilize *HostSim*, our novel mathematical and computational model that captures the dynamics of Mtb infection in a primate host at a whole-host scale. We use *HostSim* to directly investigate the cellular mechanisms that lead to concomitant immunity by relating events and dynamics within each individual virtual host to population scale outcomes. A significant benefit of this systems biology approach is that the group of 500 virtual individual *HostSim* simulations is identical across reinfection studies. Therefore, we can directly compare each reinfection study against the others. This allows us to build on the studies begun in NHP (29) and perform simulated studies impossible to do with *in vivo* experiments. Our predictions add support to growing evidence (22,29) that primary Mtb infection provides a concomitant immune response against reinfection. Further, we predict that longevity of concomitant immunity against Mtb is intrinsically tied to the magnitude of numbers of Trm cells generated, as our studies indicate that these cells are key mediators of protection against reinfection.

Trms offer intriguing targets for vaccination. In fact, intravenously administered BCG was recently shown to provide a pool of lung Trms that exhibited protection against infection with Mtb in NHPs (34). The longevity of these cells has implications for vaccine design. In humans, the longevity of Trms in the lungs is not yet known. In this work, we create a very simple model of Trms where we assume there is some level of them present in the lung after a long Mtb infection, and then allow them to decay due to a lifespan over time. This allows us to predict that in the context of an ongoing, persistent Mtb infection, the upper bound lifespan (1/*d*_*Trm*_) of Trms within the lungs is approximately 2 to 3 years. To our knowledge, this is the first estimate of Trm lifespan in human lungs and contributes to growing evidence that this long-lived cell population can persist >1 year (46). Note that this is likely an overestimate of the lifetime; it is likely that Trms continue to be generated throughout a persistent infection and thus a shorter lifetime would be sufficient to account for the data. The creation of a mechanistic mathematical model of Trms during TB together with NHP datasets is the subject of ongoing studies in our groups.

Our findings suggest that if immunity can only be achieved through an antigen-specific Trm cell population, then a successful vaccine must extend the durability, magnitude or longevity of this cell population, thereby toggling the properties of T cell memory across time. Macaque studies with a CMV vector-based TB vaccine suggest that this vaccine platform might provide the necessary antigen persistence for continuous stimulation toward a protective memory T-cell pool across time (47). However, vaccine protection studies in NHPs have yet to be completed where the timing of infection was >1 year after vaccination (47).

In the absence of Trm populations, we still note limited protection against reinfection (Figure 3D and Figure 5). In our first set of reinfection studies, the average reduction of risk of developing active TB was 35% for reinfection study numbers 2-21 (the studies that included a reinfection timepoint after the loss of all lung Trms). This percentage is consistent with estimates of protection from reinfection in a household contact study in Peru, that showed 3-35% protection against reinfection across an average interval of 3.5 years (26). Our works suggests that in addition to Trms, there are other mechanisms within the body to prevent uncontrolled bacterial growth following reinfection and perhaps this is reflected in the study of Peruvian individuals. We posit that individuals develop active TB following reinfection due to inadequate adaptive immune responses where the magnitude of both Trms and circulating Mtb-specific T cells must be insufficient.

In this study, we could not model every cell type present in both primary infection and reinfection granulomas. For example, unrestricted or unconventional T cells have not been considered in this work but could influence reinfection granuloma environments and have important implications for vaccination (12). These cell types (as well as others) could be considered in future work. Additionally, while natural TB case studies have shown protection against reinfection (22,25,26), patients who successfully complete TB drug treatment are at an increased risk of developing TB upon reinfection, with an incidence rate four times that of primary infection (48,49). We do not capture this population in the current study, although current work in our group is incorporating drug treatment into *HostSim* and could further investigate this phenomenon.

## Methods

### The HostSim modeling framework

Recently we presented *HostSim*, a whole host computational and mathematical modeling framework of Mtb infection (42). Briefly, *HostSim* tracks the development of multiple lung granulomas, as well as immune cell trafficking to the lymph node and from the blood. Each granuloma is an agent that is placed stochastically within the boundary of a 3-dimensional lung environment. Within each lung granuloma, *HostSim* captures the dynamics of various immune cells across time, including resting, infected and activated macrophages and T cells (both cytotoxic and Th1 primed, effector and effector memory cell populations), as well as pro-inflammatory and anti-inflammatory cytokines.

We additionally describe the initiation of adaptive immunity within a lymph node (LN) compartment after receiving signals from antigen presenting cells migrating from lungs. In the LN model, which has been previously presented (42,50,51), we track specific and non-specific CD4+ and CD8+ naïve, effector, effector memory, and central memory T cell responses using a compartmentalized system of 31 non-linear ordinary differential equations. Finally, we track immune cell counts within a blood compartment that acts as a bridge between lymph nodes and lungs. See Supplementary Materials for model equations, parameter values and details.

### Virtual hosts and TB outcomes

Using *HostSim*, we sample parameter space to create a virtual population of hosts and use it to delineate TB outcomes across this virtual host population. In this work, and similar to previous work (42), we identify Mtb eliminators, whose total lung CFU<1, active TB cases, whose total lung CFU >10^5^, and LTBI, all other virtual hosts following infection. Supplementary Figure S1 displays the breakdown of each clinical classification following primary infection, wherein across 500 virtual hosts, 110 are classified as active TB cases, 366 are classified as LTBI cases, and 24 are classified as Mtb eliminators.

### Including resident memory T cells during reinfection in HostSim

To test the proposed role of Trms in concomitant immunity, we expanded *HostSim* to include this cell population. To do this, we have two key assumptions about the roles of Trms during reinfection. First, Trms have been shown to kill pathogens upon re-encounter very quickly, presumably by rapidly activating macrophages to kill their intracellular bacteria (29,30,33). Therefore, we assume that Trms in *HostSim* assist infected macrophages in killing intracellular bacteria at a rate proportional to bacteria, macrophage, and Trm counts within the granuloma.

Second, Mtb-specific Trms have been identified in uninvolved lung tissue during primary infection (29). Thus, we assume Trms would be present at very early stages prior to granuloma formation upon reinfection. To identify how many Trms might be present, we compare the reinfection CFU dynamics from the NHP study (29) at the reinfection timepoint (day 112) and use 1 to 10 Trm as initial conditions (Figure 1&4). However, when we perform the parallel virtual reinfection studies (see below), we change the initial condition of the Trm population to reflect the lifespan of this cell population (see Trm lifespan section below). Based on these assumptions, we expanded the set of granuloma ordinary differential equations in *HostSim* to include an equation for Trms during reinfection scenarios (Figure 4). (See supplement for equations).

**Figure 4:**
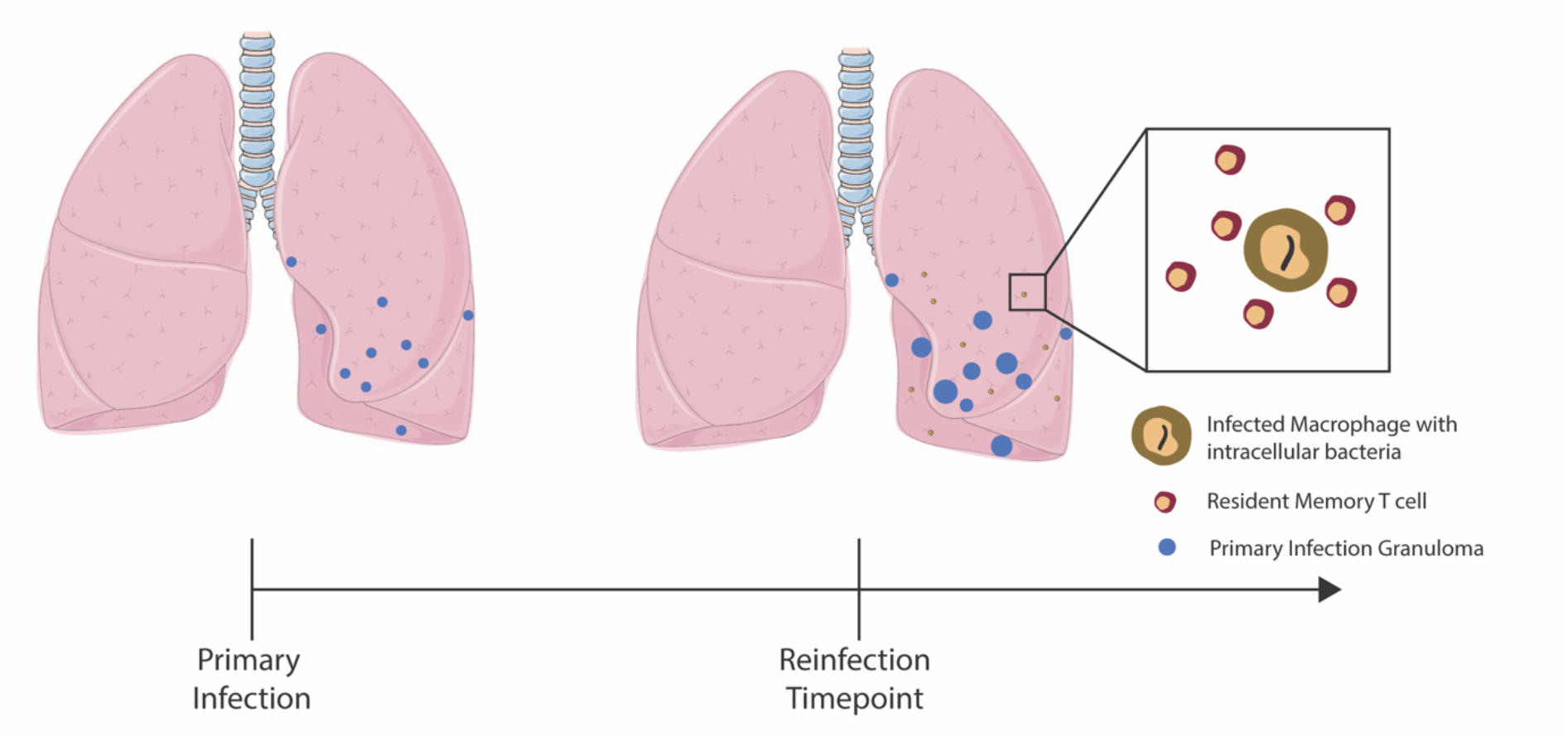
Resident Memory T cells impact establishment of reinfection. Primary infection begins with the inoculation of 10 CFU. At the reinfection timepoint, which varies from study to study, virtual hosts are re-inoculated with 10 CFU. Trm numbers vary at the time of reinfection depending on the Trm lifespan and the reinfection timepoint of the virtual study.

### Resident memory T cell (Trm) lifespan in the lungs

The lifespan of lung Trm in humans is largely unknown, but there is evidence that Trm populations are not as stable in the lung environment as they are in the skin or other locations (52,53). Morris et al used an exponential decay function to model the longevity of Trm populations across time and assumed no influx to the lung Trm population:

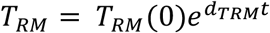

where *d*_*DRM*_ is the death rate of Trm (43). By calibrating to mice lung Trm datasets, Morris et al. determined *d*_*TRM*_ = 0.03 cells/day, or a lifespan of about 33 days (where lifespan is calculated as 1/*d*_*Trm*_). As they note, human lung Trms are less well-studied, primarily due to inability to sample healthy lung tissue. In our parallel virtual host reinfection studies (see below), we utilize this Trm longevity function to inform the initial conditions of Trms in reinfection scenarios. Additionally, we test different values for Trm death rates to predict the lifespan of these cell types in primates.

### Reinfection events in HostSim

Reinfection studies within *HostSim* involve re-inoculating our virtual hosts with 10 CFU at the reinfection timepoint, which seeds 10 unique new granulomas within the lung environment (Figure 4). Each reinfection granuloma begins with one intracellular bacterium, one infected macrophage and between 1 and 10 Trms as initial conditions.

### Parallel virtual host reinfection studies

To predict the lifespan of Trms in the lung, we perform several parallel virtual host studies. Figure 5 demonstrates the experimental protocol for three sets of 21 reinfection studies. Each reinfection study consists of the same 500 individual *HostSim* simulations run for 500 virtual hosts over approximately a 7-year timeframe (2,500 total days) differing only in the time of reinfection. Study 1 begins with a reinfection timepoint at day 112 (the reinfection timepoint used in the NHP dataset from Cadena et al. (29)) and each subsequent study has a reinfection time point occurring 100 days after the previous study.

**Figure 5:**
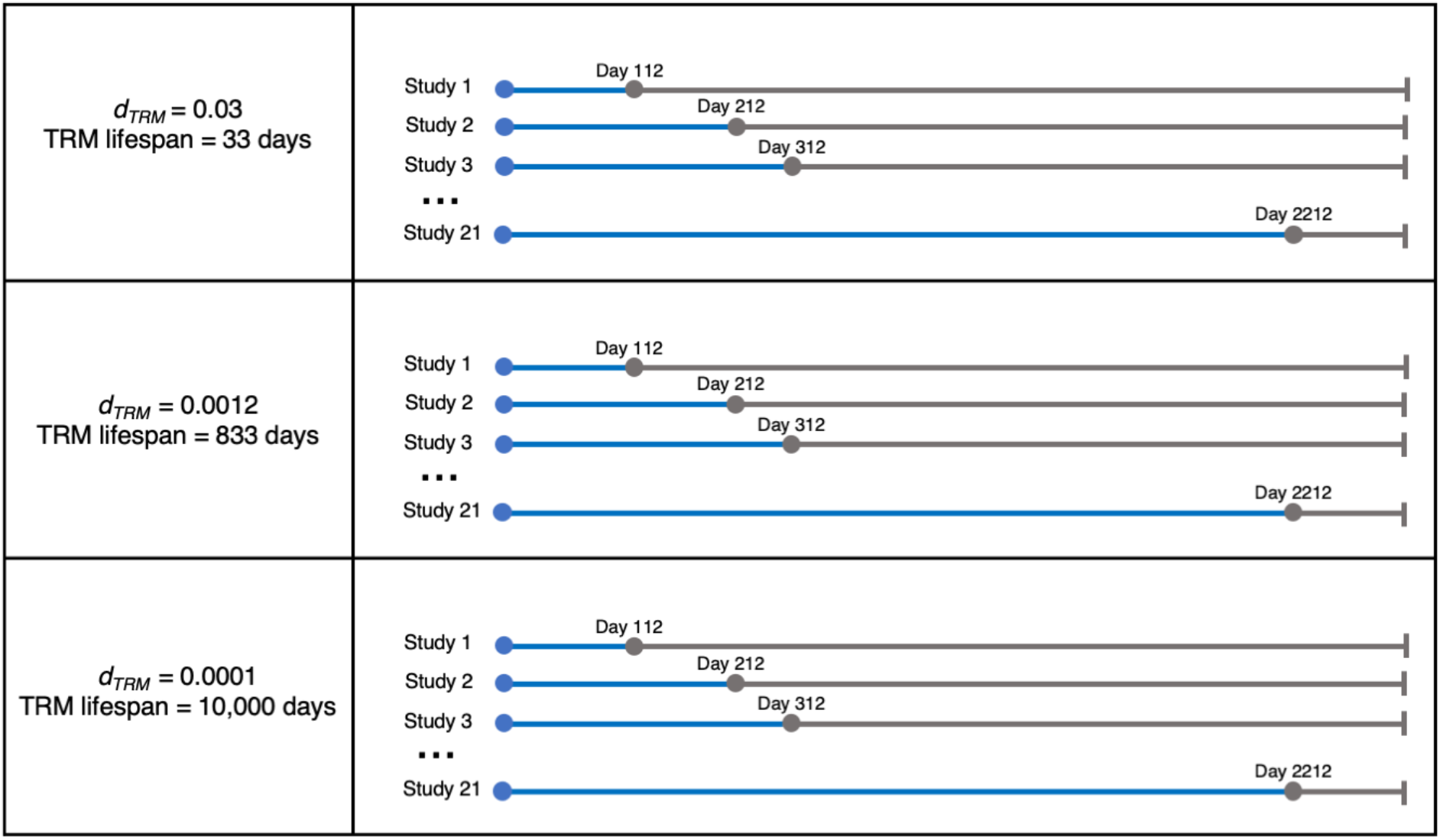
Parallel virtual host reinfection studies to predict Trm lifespan. The experimental protocol for our *HostSim* reinfection studies. We performed 3 sets of 21 separate reinfection studies on 500 virtual hosts, where the only difference between studies was timing of Mtb reinfection with a low dose of 10 CFU. Reinfection time points shown as gray dots on the timeline. We assigned a different death rates of Trm (*d*_*Trm*_) for each of 3 sets of reinfection studies. In total, we perform 63 reinfection studies where the same 500 hosts are simulated for 2500 days (approx. 7 years).

The 21 reinfection studies are each simulated three times, varying the death rate of Trms between each set. The first death rate, 0.03 cells/day, is the predicted death rate of Trm in mice from Morris et al (43). The second, 0.0012 cells/day, was calculated based on allometric scaling, as we assume humans live ~25x longer than mice (54). The third death rate selected was 0.0001 cells/day, a rate similar to that observed for central memory T cell populations, which are known to persist for decades (55).

### Calculating reduced risk of active TB following reinfection compared to primary infection

Studies from the pre-antibiotic era of TB tracked the percentage of nursing students that contracted active TB disease (22). Upon entering nursing school, students were stratified into two groups: those that had a positive Tuberculin Skin Test (TST), indicating previous exposure to Mtb, and those that had a negative TST, indicating no previous exposure to Mtb. A meta-analysis of these studies showed that individuals with a TST+ upon entering nursing school were at a 79% reduced risk of developing active TB compared to TST-individuals (22). Assuming that all nurses had similar exposure to infectious TB patients, these studies are strong evidence of a protective concomitant immune response against reinfection. Supplementary Figure S2 shows the data from these studies – note that studies with a longer period of observation (closer to 5 years) saw greater percentages of active TB cases among the TST+ population compared to those where students were observed for two years or less. Although this association is not statistically significant, the greater percentage of active TB cases during longer studies suggests a potentially waning protection from concomitant immunity.

Using *HostSim*, we can estimate the reduced risk of developing active TB from reinfection for virtual hosts. We do this by simulating a virtual population of hosts and comparing 1) the fraction of virtual hosts which develop active TB from primary infection and 2) the fraction of virtual hosts which develop active TB from reinfection. These two fractions are analogous to the TST- and TST+ groups in the meta-analysis of nursing students, respectively (22). As Supplementary Figure S1 shows, 110 out of a virtual population 500 hosts will develop active TB (total lung CFU>10^5^ at day 200 post-infection) following primary infection. This number (110/500) acts as our expected value following infection. We determine the second fraction during our reinfection studies by counting the number of hosts that have a total lung CFU > 10^5^ at 200 days after reinfection. As an example, if 50 out of 500 virtual hosts develop active TB after reinfection, the risk reduction calculation would be as follows (in absolute value):

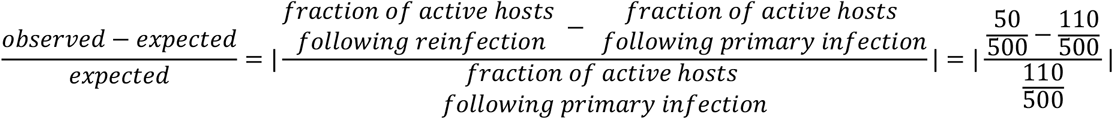

resulting in a 54.5% reduced risk of developing active TB.

### Model environment and analysis

Model code and preliminary data analysis is written in MATLAB (R2020b). ODEs are solved using MATLAB’s ode15s stiff solver. Simulation for a single *in silico* individual across 200 days post-infection can be performed on an 8-core laptop in approximately 30 seconds. Bash scripts were written for submission to run on the Great Lakes HPC Cluster at the University of Michigan for parallel virtual host reinfection studies. Post-processing statistical analysis and graphing was performed in MATLAB (R2020b).

## Acknowledgements

This research was supported by NIH Grants R01 AI50684 (DEK JLF) and U01 HL131072 (DEK, JJL). Additionally, this work is supported by funding by the Wellcome Leap HOPE Program awarded to (DEK, JJL, JLF). LRJ was funded by a University of Michigan Rackham Predoctoral Fellowship. Simulations also use resources of the National Energy Research Scientific Computing Center, which is supported by the Office of Science of the U.S. Department of Energy under Contract No. ACI-1053575 and the Extreme Science and Engineering Discovery Environment (XSEDE), which is supported by National Science Foundation Grant MCB140228.

