## Supplementary Materials for "Concomitant immunity to M. tuberculosis infection"

### Supplementary Material

*HostSim Equations with the addition of Resident Memory T cells*

The *HostSim* granuloma model equations are shown in full detail in previously published work (42). Briefly, the system of 20 ordinary differential equations captures intracellular and extracellular bacterial numbers, various macrophages, various T cells and pro- and anti-inflammatory cytokines across time. In this work, we add a resident memory T cell population (Trm) to reinfection granulomas:

$$T_{rm} = T_{rm}(0)e^{d_{Trm}t}$$

where $d_{TRM}$ is the death rate of Trm and is assigned as 0.03, 0.0012, or 0.0001 cells/day, depending on the study. $T_{rm}(0)$ = [1-10] and is sampled according to a Latin Hypercube sampling scheme (56), like other parameters in *HostSim*. All other equations remain unchanged, except intracellular bacteria, which now includes a term from interactions between macrophages and Trm cells that leads to intracellular bacterial death:

$$\frac{dB_I}{dt} = \alpha_{19}\frac{B_I}{M_I}M_I\left(1-\frac{\frac{B_I}{M_I}}{N}\right) + k_2\frac{N}{2}M_R\left(\frac{B_E}{B_E+c_9}\right) - k_{17}NM_I\left(\frac{B_I^2}{B_I^2+N^2M_I^2}\right) - k_{14a}\frac{B_I}{M_I}M_I\left(\frac{\left(\frac{T_C+w_3T_1}{M_I}\right)}{\left(\frac{T_C+w_3T_1}{M_I}\right)+c_4}\right)$$

$$- k_{14b}\frac{B_I}{M_I}M_I\left(\frac{F_\alpha}{F_\alpha+f_9I_{10}+s_{4b}}\right) - k_{52}\frac{B_I}{M_I}M_I\left(\frac{\left(\frac{\frac{T_C}{B_i+1}\left(\frac{T_1}{T_1+c_{T_1}}\right)+w_1T_1}{M_I}\right)}{\left(\frac{\frac{T_C}{B_i+1}\left(\frac{T_1}{T_1+c_{T_1}}\right)+w_1T_1}{M_I}\right)+c_{52}}\right) - \mu_{B_I}B_I + \mu_{M_I}\frac{B_I}{M_I}M_I$$

$$- k_{BIdeathTRM} * M_I * T_{rm} * \frac{B_I}{M_I}$$

Where $k_{BIdeathTRM}$ = [0.3, 0.8] bacteria/day consistent with rate constants of intracellular bacteria death from interactions with other T cells in *HostSim*, and identified via manual tuning to match the reinfection CFU dynamics of the NHP study by Cadena et al (29).

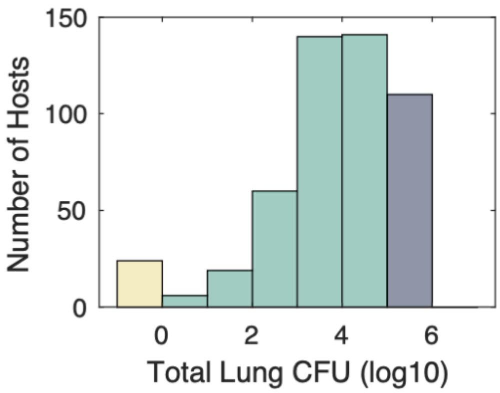


**Supplementary Figure S1: Histogram of Total Lung CFU in *HostSim* virtual population of 500 hosts.** Total lung CFU calculated by summing CFU across all granulomas in a host at day 200 following primary infection. Yellow represents hosts that are classified as Mtb eliminators (total Lung CFU < 1), dark blue represents hosts classified as active TB cases (total Lung CFU > $10^5$) and green represents latently infected individuals (LTBI), i.e. those that control infection. Across a population of 500 hosts, 110 are classified as active TB cases, 366 are classified as LTBI, and 24 are classified as Mtb eliminators. This breakdown of responses across a virtual population of 500 individuals represents our expected outcomes when calculating reduced risks of developing active TB following reinfection.

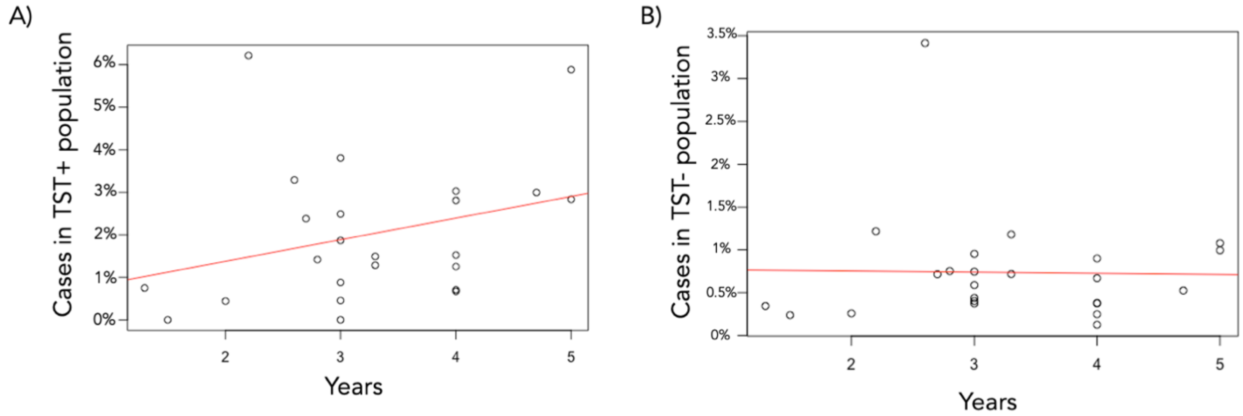


**Supplementary Figure S2: Percentages of active TB cases per study for TST+/TST- individuals prior to the study.** A) The percentage of TST+ individuals who develop active TB across time in the meta-analysis by Andrews et al. (22). B) the percentage of TST- individuals who develop active TB across time. Each data point is a single study, where the x-axis represents the length of the study.

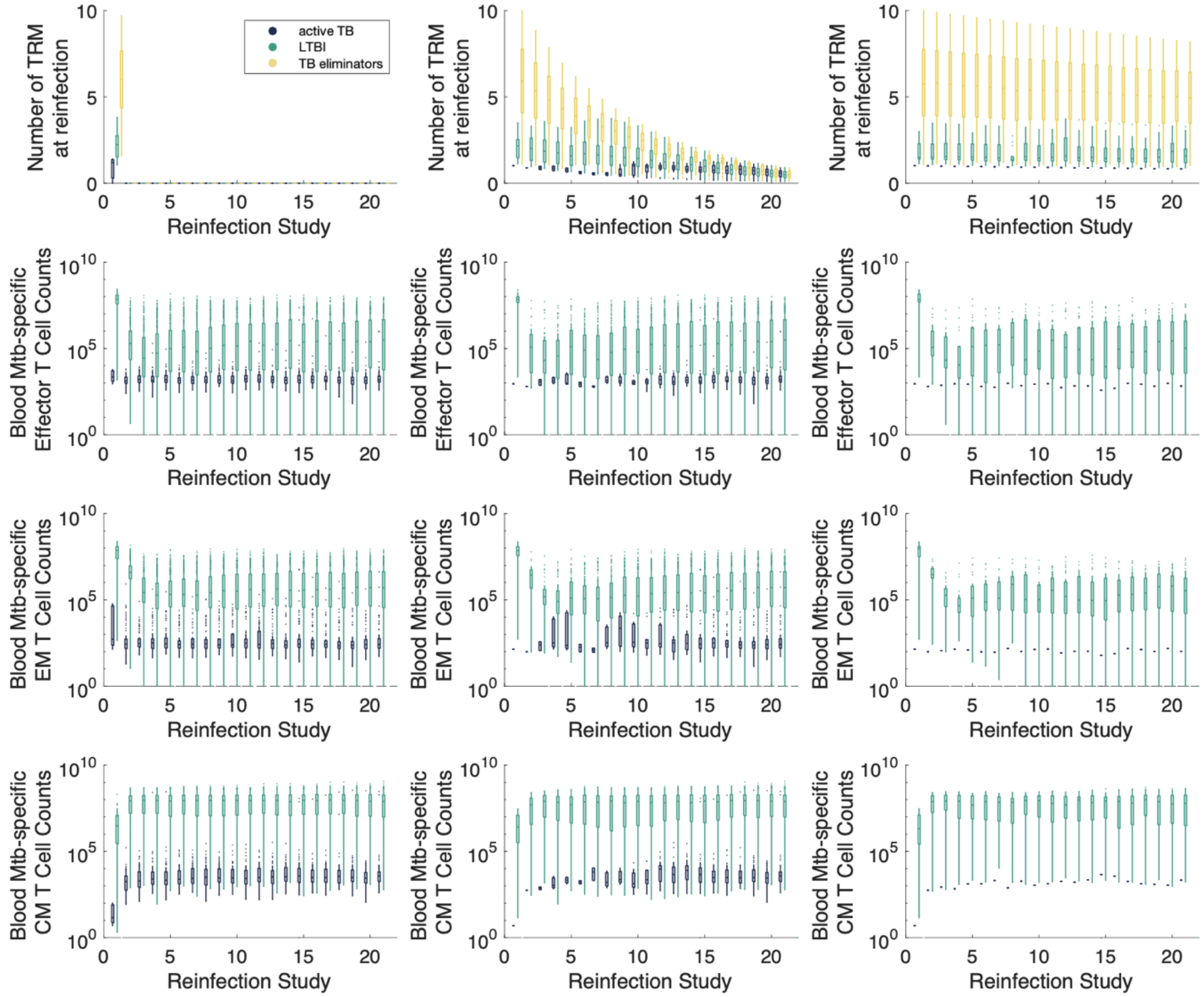


**Supplementary Figure S3: Blood T cell counts delineate active TB and LTBI cases for each set of reinfection studies.** Box-and-whisker plots show the distribution of Trm, Mtb-specific effector, effector memory or central memory T cells in the blood for hosts that were active TB, LTBI or Mtb eliminators following reinfection. Dark blue=active TB, Green = LTBI, yellow=Mtb eliminator. Each column represents each set of the three reinfection studies; where the death rate of Trm cells from left to right is $d_{Trm}$ =0.03, $d_{Trm}$ =0.0012, and $d_{Trm}$ =0.0001 cells/day.
